# Y-RNAs Lead an Endogenous Program of RIG-I Agonism Mobilized upon RNA Virus Infection and Targeted by HIV

**DOI:** 10.1101/773820

**Authors:** Nicolas Vabret, Valérie Najburg, Alexander Solovyov, Petr Šulc, Sreekumar Balan, Guillaume Beauclair, Maxime Chazal, Hugo Varet, Rachel Legendre, Odile Sismeiro, Raul Y. Sanchez David, Christopher McClain, Ramya Gopal, Lise Chauveau, Olivier Schwartz, Nolwenn Jouvenet, Martin Markowitz, Frédéric Tangy, Nina Bhardwaj, Benjamin D. Greenbaum, Anastasia V. Komarova

**Affiliations:** Tisch Cancer Institute, Icahn School of Medicine at Mount Sinai, New York, New York, USA.; Precision Immunology Institute, Icahn School of Medicine at Mount Sinai, New York, NY, 10029, USA; Viral Genomics and Vaccination Unit, Department of Virology, Institut Pasteur, CNRS UMR-3569, 75015 Paris, France; Center for Molecular Design and Biomimetics at the Biodesign Institute and School of Molecular Sciences, Arizona State University, Tempe, Arizona 85287, USA; Transcriptome and EpiGenome platform, BioMics, Center of Innovation and Technological Research, Institut Pasteur, 28 rue du Docteur Roux, 75724 Paris Cedex 15, France; Hub informatique et Biostatistique, Centre de Bioinformatique, Biostatistique et Biologie Intégrative (C3BI, USR 3756 IP-CNRS), Institut Pasteur, 28 Rue du Docteur Roux, 75724 Paris Cedex 15, France; Virus & Immunity Unit, Department of Virology, Institut Pasteur, CNRS UMR-3569, 75015 Paris, France; Aaron Diamond AIDS Research Center, The Rockefeller University, New York, NY, USA; Parker Institute of Cancer Immunotherapy, USA; Department of Medicine, Hematology and Medical Oncology, Department of Oncological Sciences, Department of Pathology, Center for Computational Immunology, Tisch Cancer Institute, Icahn School of Medicine at Mount Sinai, New York, NY, 10029, USA; Icahn Institute for Data Science and Genomic Technology, Icahn School of Medicine at Mount Sinai, New York, NY, 10029, USA.; Viral Genomics and Vaccination Unit, Department of Virology, Institut Pasteur, CNRS UMR-3569, 75015 Paris, France.

## Abstract

Pattern recognition receptors (PRRs) protect against host invasion by detecting specific molecular patterns found in pathogens and initiating an immune response. While microbial-derived PRR ligands have been extensively characterized, the contribution and relevance of endogenous ligands to PRR activation during viral infection remain overlooked. In this work, we characterize the landscape of endogenous ligands that engage RIG-I-like receptors (RLRs) upon infection by a positive-sense RNA virus, a negative-sense RNA virus or a retrovirus. We found that several endogenous RNAs transcribed by RNA polymerase 3 (Pol3) specifically engage RLRs, and in particular the family of small non-coding repeats Y-RNAs, which presents the highest affinity as RIG-I ligands. We show that this recognition is dependent on Y-RNA mimicking viral secondary structure and its 5’-triphosphate extremity. Further, we found that HIV-1 infection triggers a VPR-dependent downregulation of RNA triphosphatase DUSP11 *in vitro* and *in vivo*, leading to an increase of Y-RNA 5’-triphosphorylation that enables their immunogenicity. Importantly, we show that altering DUSP11 expression is sufficient to induce a type-I interferon and T cell activation transcriptional program associated with HIV-1 infection. Overall, our work uncovers the critical contribution of endogenous repeat RNAs ligands to antiviral immunity and demonstrates the role of this pathway in HIV-1 infection.

## Main

Pattern Recognition Receptors (PRRs) were initially described as innate immune sensors of molecular patterns commonly found in pathogens but rarely, if ever, found in their hosts. In recent years, this view has been challenged by evidence that ligands originating from self can engage these same PRRs. Notably, sensing of self-RNA by innate receptors has been observed in various settings such as autoimmune disorders (*1-3*), tumorigenesis and cancer therapies (*4-9*) or infection by DNA viruses (*10, 11*). While the importance of endogenous ligands in priming immune responses is progressively uncovered, little is known about the breadth of biological processes in which they happen, nor about their functional and evolutionary interplay with immune sensors. Furthermore, we lack understanding of what features confers self-RNAs the ability to activate sensors and whether this is a general response to aberrant transcription or is dominated by specific RNA species.

Further confounding matters, we previously determined that conventional RNA sequencing approaches fail to capture the full spectrum of RNA expression in tumors (*12*). In particular, repetitive RNA, which can harbor immunostimulatory features (*13*), require further computational analysis for unbiased screening of their transcription. Here, we apply these approaches to identify novel RNA agonists of RIG-I-like receptors (RLRs). RLRs are a family of cytosolic RNA sensors composed of three members: RIG-I, LGP2 and MDA5 (*14*). Their intracellular localization and proximity with host RNA species implies a delicate balance between a need to develop high affinity for microbial features and the possibility to encounter self-RNAs that display similar structures. However, despite a growing knowledge of the role of RLRs during RNA virus infection and the microbial-derived ligands they recognize (*14*), the contribution of specific endogenous RNAs to their activation and the mechanisms controlling their immunogenicity remain elusive.

### Y-RNAs and other RNA Pol3 transcripts are cellular RIG-I ligands mobilized upon RNA virus infection

We recently developed a riboproteomic approach based on tagged protein affinity purification that measures and compares receptor affinity of RNA molecules with improved statistical evaluation of specific binding (*15, 16*). We performed an unbiased quantification of RLR-bound self RNAs during RNA virus infection. We generated human HEK293 (293) cells stably expressing the tagged-RLRs RIG-I, MDA5 or LGP2, or the protein Cherry as non-binding control. We infected each cell line with either positive-sense RNA virus Dengue Virus 4 (DV-4) or negative-sense RNA virus Measles Virus (MV). As a model of retroviral infection, we co-cultivated HIV-1-infected MT4 T cells with 293 cells overexpressing HIV-1 receptors CD4 and CXCR4 (293-4×4), as cell-free HIV-1 particles are poor stimulators of type I interferon (IFN-I) (*17*). We performed total RNA-sequencing on each RLR-or Cherry-purified fraction and on total cellular RNA (Fig. S1A). Importantly, we confirmed that the RLR-MAVS pathway is critical for sensing each viral infection in this model (Fig. 1A, Fig. S1B). We previously reported specific viral RNA-binding profiles on RLR compared to non-specific binding (Cherry) upon MV and DV-4 infections (*15, 16*) (Fig. S1C-D). However, upon HIV-1 infection, no enrichment of viral RNA was observed on any receptors (Fig. S1E). We then aligned RLR-bound RNAs to the human genome and measured specific cellular RNA enrichments in infected and non-infected conditions. Importantly, we found a strong enrichment of Pol3-transcribed RNAs to RIG-I and LGP2 upon each RNA virus infection, and in particular Y-RNAs (Fig. 1B-C, Fig. S1F, S1H, Table S1). Y-RNAs constitute a family of highly conserved small noncoding RNAs transcribed by Pol3, composed of four canonical Y-RNA (RNY1, RNY3-5) and several hundreds of pseudogenes (*18*). As the repetitive nature of Y-RNAs makes it impossible to identify the exact origin of each transcript, we measured RLR enrichment of each repeat family rather than individual genes (Fig. 1D, S1G, S1I, Table S2). Specifically, the subfamily of HY4, which contains RNY4 and its pseudogenes, and to a lesser extent HY3, showed significant binding enrichment to RIG-I in the three RNA virus infections compared to non-infected conditions (NI).

**Figure 1:**
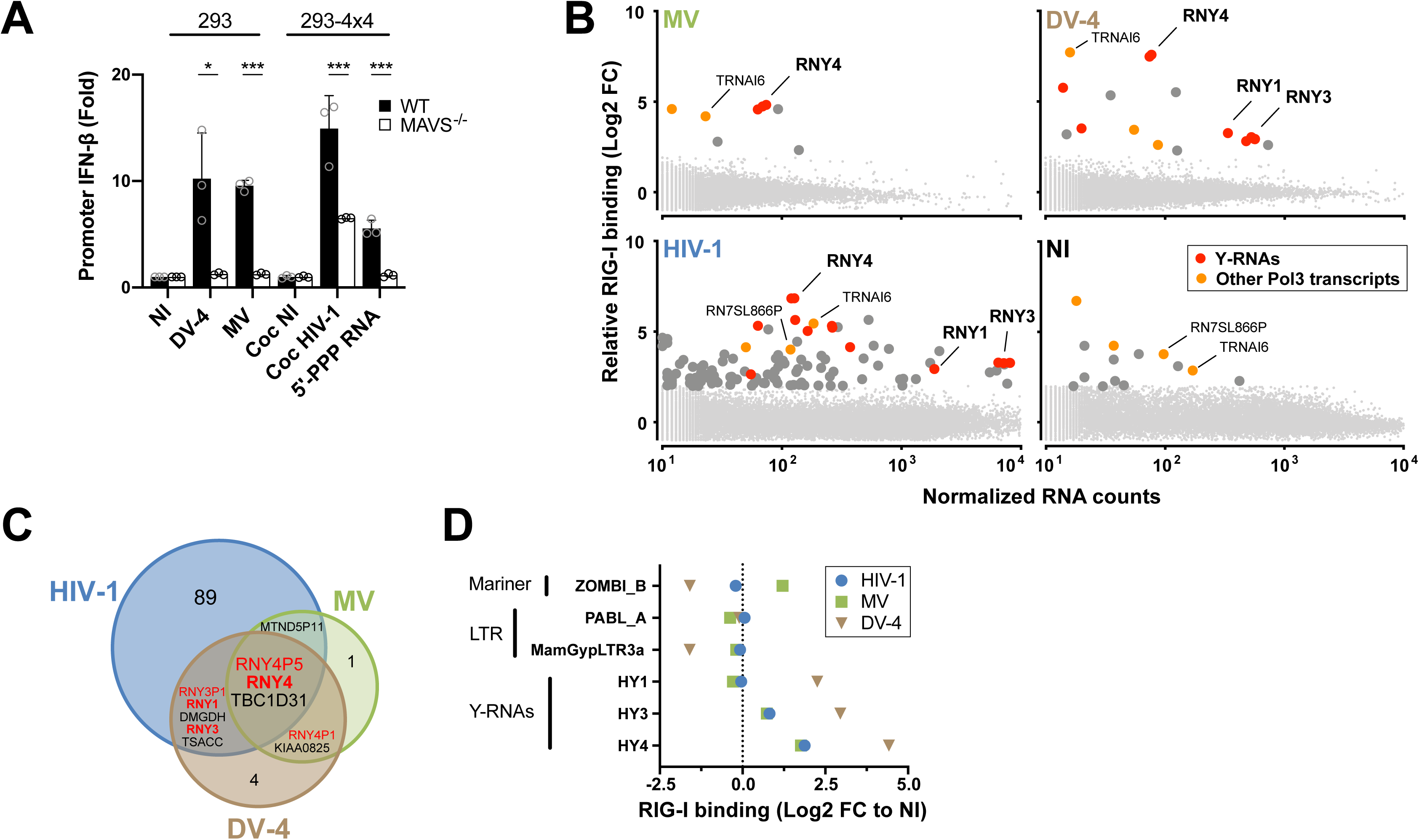
A differential affinity screen identifies Y-RNAs and other POL3 RNAs as RIG-I ligands mobilizable during RNA virus infection. **A.** Promoter IFN-β-luciferase reporter activity in WT or MAVS-/- (left) 293 cells infected with Measles virus (MV) or Dengue Virus 4 (DV-4) at MOI of 1 and 0.5 respectively, or (right) 293-4×4 cocultivated with HIV-1 infected MT4C5 at a ratio of MT4C5:293-4×4 of 1:1. 5’-PPP is a short *in vitro* transcribed RNA RIG-I agonist transfected at a concentration of 10 ng/ml. **B.** 24h post-infection with MV or DV-4, after coculture with HIV-1 infected MT4 or in non-infected (NI) control, sequencing reads were mapped to human genome Hg38. Differential analyses were performed between RIG-I/RNA and Cherry/RNA samples. Genes are represented following their normalized count in cellular RNA (x-axis) and their fold enrichment (log2) to RIG-I compared to Cherry control (y-axis) from averaging three independent replicates. Genes that showed a log2(FC)>2 and adj-pval<0.05 are represented with larger dot size. Among these, Pol3 transcripts are shown in orange and transcripts from Y-RNA families in red. Canonical Y-RNAs and Pol3 transcripts that show enrichment in more than 2 conditions (virus or NI) are specifically annotated. **C.** Venn diagram representing genes specifically enriched to RIG-I compared to Cherry in any of MV, DV-4 or HIV-1 infected conditions, but absent in NI condition. **D.** Families of repeats RNA that show specific affinity to RIG-I compared to Cherry in at least one infected or NI condition, computed according to their relative enrichment compared to NI. **A.** Data representative of n=3 independent experiments. Bars show mean +/− s.e.m. of technical triplicates. Student’s t-test *p<0.05; ***p< 0.001. **B-D.** Enrichment calculated from the mean of n=3 infection/RLR-purification/sequencing experiments.

### 5’-PPP and a specific secondary structure are required for RNY4 RIG-I agonist activity

To analyze the immunostimulatory properties of Y-RNAs, we generated *in vitro* transcribed (IVT) molecules of each canonical Y-RNA and measured IFN-I signaling after stimulation of individual RLR knock-outs generated in the haploid cell line HAP1. Each individual Y-RNA was able to elicit an IFN-I response after transfection, which was dependent on the presence of RIG-I and MAVS, but independent of MDA5 or to a large extent LGP2 (Fig. 2A). Further, as cellular Y-RNAs are observed under two different forms, a full-length form and shorter fragments derived from its 5′ and 3′ termini (*18*), we compared RNY4 reads coverage between the fraction bound to RIG-I and the fraction sequenced from the total cellular RNA pool in the different experimental infections settings. Interestingly, we did not find significant differences between RIG-I-bound RNAs and total cellular RNY4, and coverage results suggested that the uncleaved form of RNY4 controls its RIG-I binding property (Fig. S2A). RIG-I recognizes RNA ligands based on a level of specificity in terms of sequence composition, length, double-stranded structures, and presence of triphosphate (-PPP) or diphosphate 5’ moieties(*14*). We generated fragments derived from RNY4 missing specific molecular substructures (Fig. S2B). As shown in Fig. 2B, both 5’-PPP and stem S3 were required to confer upon RNY4 its RIG-I-dependent immunostimulatory activity.

**Figure 2.**
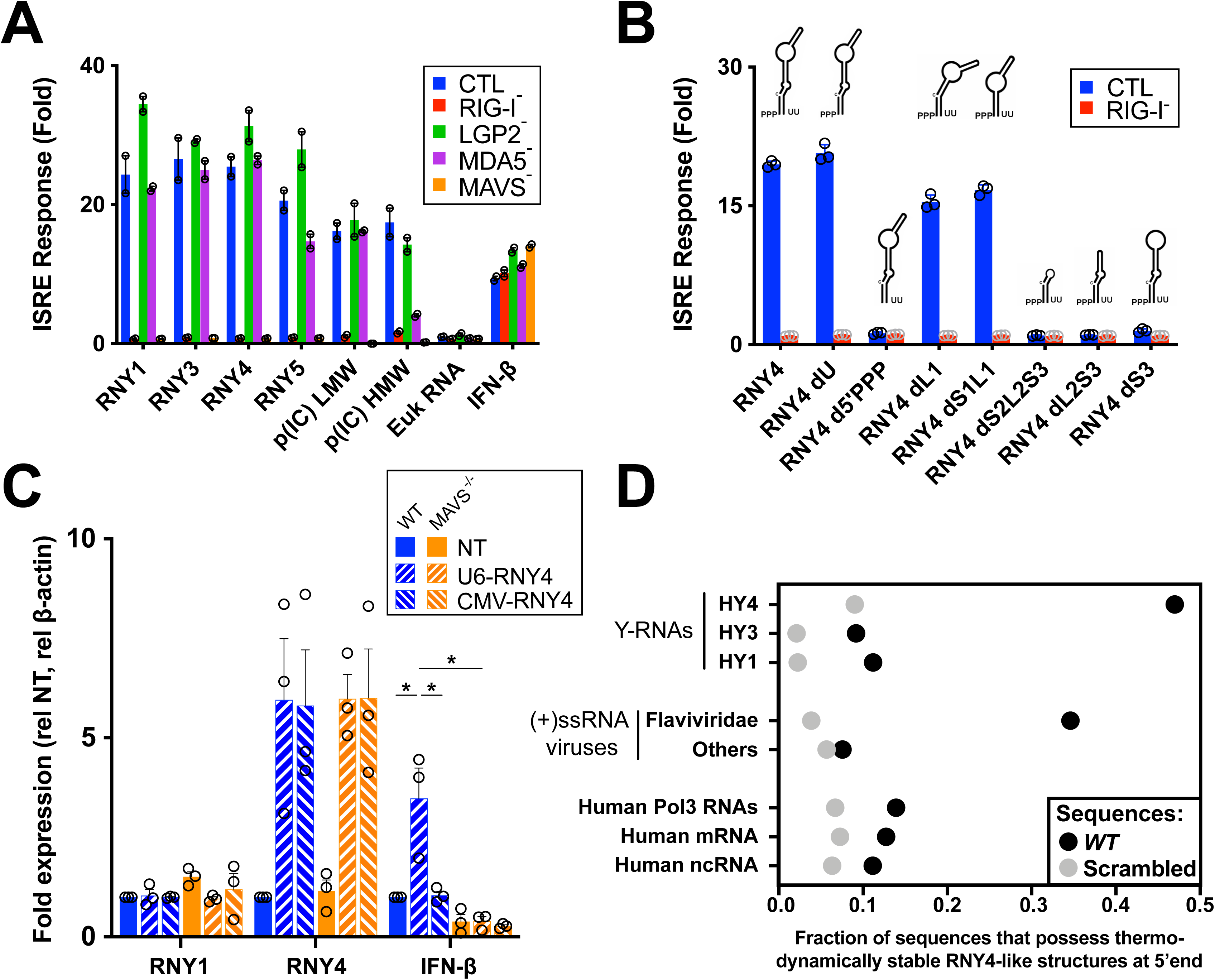
RNY4 RIG-I agonist activity is conferred by RNA 5’-PPP moieties and viral-mimicking specific secondary structure. **A.** Promoter ISRE-luciferase reporter activity in HAP1 cells WT or ko for each individual RLR or downstream adaptor MAVS, transfected with 30ng/ml of IVT Y-RNA (RNY1, RNY3-5), 10ng/ml poly(I:C) low or high molecular weight (p(IC) LMW/HMW), 30ng/ml of total eukaryotic RNA or treated with 100U/ml recombinant IFN-β. **B.** Promoter ISRE-luciferase reporter activity in HAP1 cells WT or RIG-I ko transfected with 30ng/ml IVT RNY4 full length or lacking specific substructure (Fig. S2B). RNY4 d5’PPP: RNY4 was additionally pretreated with alkaline phosphatase to remove 5’ triphosphate extremity. **C.** RNY1, RNY4 and IFN-β RNA levels measured by qPCR after transfection of 293T *WT* or MAVS^-/-^ with plasmids coding for RNY4 sequence and supplemented with a plasmid coding for RIG-I. U6-RNY4: p2RZ plasmid encoding full length RNY4 downstream of Pol3 U6 promoter with 3’ ribozyme sequence. CMV-RNY4: same plasmid with Pol2 CMV promoter instead of U6. NT: empty plasmid. **D.** Probability of sequence folding along RNY4 secondary structure in the 5’ end of each transcript, for dataset of human Y-RNAs repeat families, (+)ssRNA viruses genomes (*Flaviviridae* or non-*Flaviviridae*), or human non coding RNA (ncRNA), mRNAs and Pol3 transcripts, compared to average probability of the same sequences randomly scrambled. **A-B.** Data representative of n=3 independent experiments. Bars show mean +/− s.e.m. of technical duplicates. **C.** Bars show mean +/− s.e.m of n=3 independent experiments. Student’s t-test *p<0.05.

To validate these findings, we synthesized IVT RNY4 and RNY4ΔS3 RNAs using plasmids containing 3’ ribozyme sequences that generate discrete 3’ ends. We measured IFN-I response after transfection with these RNAs, confirming the difference observed earlier between RNY4 and RNY4ΔS3 (Fig. S2C). Finally, to confirm that endogenously transcribed Y-RNAs can be immunostimulatory, we cloned the RNY4 sequence downstream of an RNA Pol3 promoter (U6) and used a Pol2 promoter (CMV) as control. Only endogenous transcription of RNY4 driven by RNA Pol3, but not when driven by Pol2, elicited an IFN-I response dependent on the RIG-I/MAVS pathway (Fig. 2C). To further understand the novel function of Y-RNAs as a RIG-I agonist, we created a secondary structure model of RNY4 and computed the probability of different sets of sequences of viral or human origins to fold along this model. Strikingly, RNY4-like structures were more often predicted in the 5’ end of positive-sense RNA virus genomic sequences than in human RNA families, and specifically in sequences from *Flaviviridae* virus family (Fig. 2D). Altogether, these results suggest that Y-RNAs from the subfamily HY4 display patterns of endogenous viral mimicry and can be mobilized as RIG-I agonists upon infection.

### DUSP11 modulates RNY4 5’-PPP and is downregulated by HIV-1 VPR

As Y-RNAs are readily expressed at steady-state, we questioned what triggers their immunogenicity upon viral infection. Our results indicate that a 5’-PPP end is required for RNY4 RIG-I agonist activity. We performed a differential enzymatic digestion assay (Fig. S3A) to analyze the 5’ structure of RNAs in HIV-1-infected Jurkat T cells. Surprisingly, HIV-1 infection induced a hyper triphosphorylation of RNY1 and RNY4 compared to non-infected cells (Fig. 3A). Every Pol3-transcribed RNA initially contains a 5’-PPP upon transcription that can be further edited by different cellular enzymes. Among these, DUSP11 is a protein from the dual-specificity phosphatase family that displays 5’-triphophatase activity on several miRNA and other cellular noncoding RNAs (*19, 20*). We generated DUSP11 knock-out Jurkat cells to analyze its activity on Y-RNAs. Deletion of DUSP11 led to a notable increase in 5’-PPP levels at RNY4 5’-end compared to WT cells, as previous results suggested (*20*) (Fig. S3B, S3C). Further, infection with an HIV-1 NL4.3 clone coding for GFP (HIV-GFP) led to profound DUSP11 downregulation in Jurkat and primary CD4 T cells (Fig. 3B-C). The predominantly nuclear localization of DUSP11 (*21*) and the rapid downregulation kinetics observed upon HIV-1 infection led us to hypothesize that HIV-1 viral protein R (VPR) could be responsible for the observed effect on DUSP11 levels. *VPR* codes for a conserved accessory protein that incorporates into viral particles, has nuclear transport ability and induces the proteasomal degradation of several host cell factors (*22*). We compared DUSP11 downregulation in Jurkat T cells infected by either *WT* HIV-1 or the same clone lacking VPR (HIVΔVPR). We found that VPR expression was required for HIV-1-induced DUSP11 downregulation (Fig. 3D). Concordantly, expression of *WT* VPR after transduction by lentiviral vectors, but not of a VPR(Q65R) mutant unable to recruit the DCAF1/DDB/Cul4 ligase complex (*23*), was sufficient to induce DUSP11 downregulation in Jurkat (Fig. S3D).

**Figure 3:**
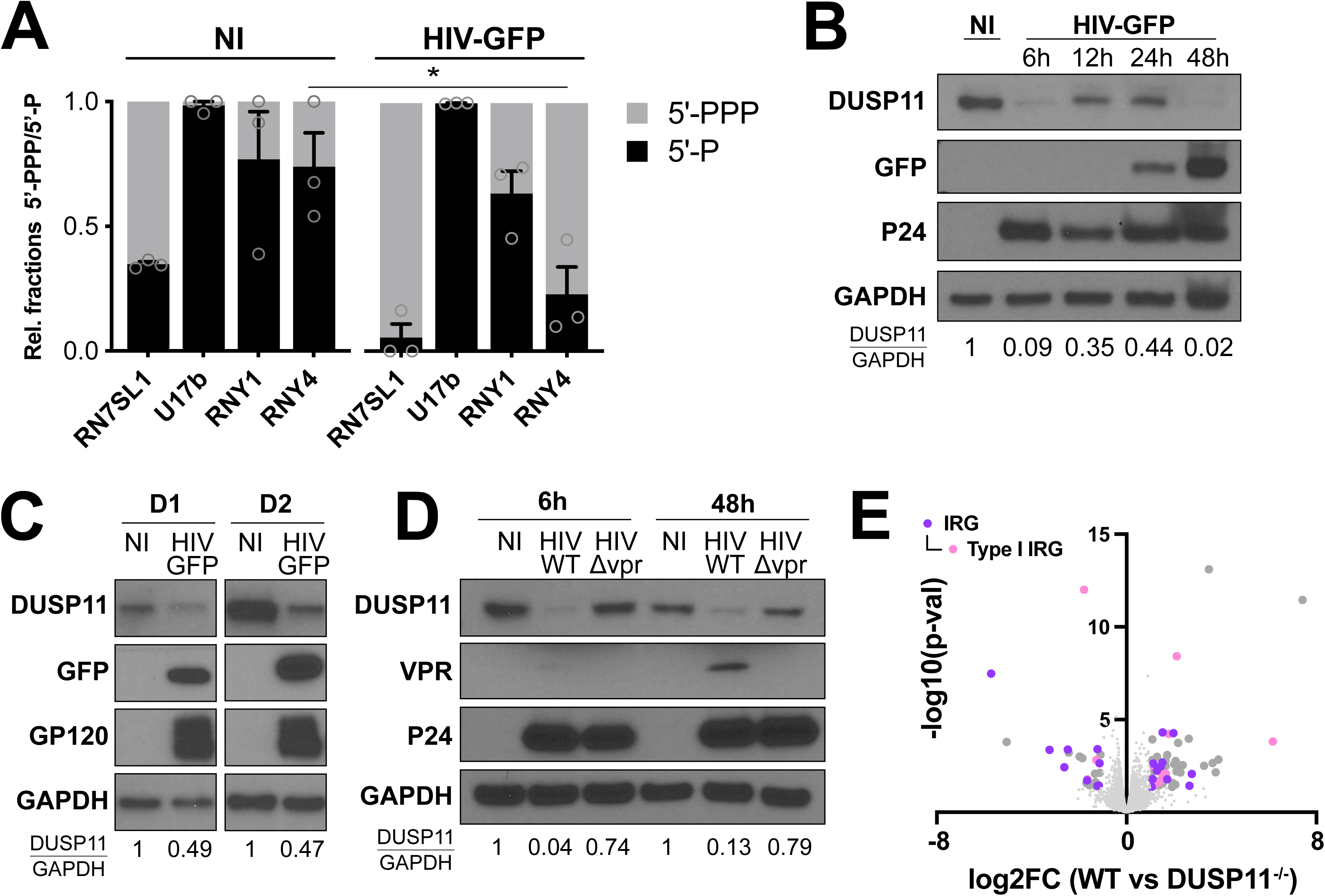
HIV-1 VPR-dependent downregulation of DUSP11 licenses endogenous Pol3-transcribed RNAs immunogenicity in infected cells. **A.** Ratio of 5’-PPP and 5’-P-bearing RNY1 and RNY4 in Jurkat cells 48h post infection with HIV-GFP or in non-infected (NI) Jurkat. Relative 5’-PPP/5’-P RNA levels were determined through differential enzyme digestion followed by qPCR analysis relative to *β-actin* mRNA. RN7SL1 and U17b are 5’-PPP and 5’-P RNA controls, respectively. **B.** DUSP11 protein levels measured at different times points after Jurkat T cells infection with HIV-GFP. **C.** DUSP11 protein levels measured in NI or HIV-GFP-infected CD4 primary cells from 2 different donors 48h post-infection. CD4 T cells were beads-sorted from total PBMC and activated with PHA for 72h prior to infection with HIV-GFP. 48h post-infection, productively infected cells were FACS-sorted according to GFP expression. **D.** DUSP11 protein levels measured at 6h and 48h after Jurkat T cells infection with *WT* NL4.3 HIV-1 or the same clone deleted for VPR protein. **E.** Volcano-plot of differential expressed genes in 3 *WT* or DUSP11^-/-^ Jurkat clones. IFN or IFN-I regulated genes (IRG) (annotation according to interferome database) are labelled in purple (dark and light, respectively). **A.** Bars show mean +/− s.e.m. of n=3 independent experiments. Student’s t-test *p<0.05. **B-D**. Western Blot representative of n=3 independent experiments. **B-D** Numbers at the bottom indicate semi-quantification of relative DUSP11/GAPDH levels normalized to NI conditions.

Since DUSP11 dephosphorylates the 5’ end of putative endogenous RIG-I ligands, we analyzed whether DUSP11 deficiency was sufficient to trigger an innate immune response by performing total RNAseq on *WT* and DUSP11^-/-^ Jurkat clones. A large fraction of differentially expressed genes were interferon-regulated genes, as annotated in the interferome database (*24*) (Fig. 3E). We further validated by qPCR the upregulation of genes from a panel of classical type-I ISGs (Fig. S3E). Additionally, we confirmed the upregulation of mRNAs coding for T cell surface markers involved in T cell activation and survival such as CD28, CD38 or IL7R that we found differentially expressed by RNAseq (Fig. S3E). Finally, we defined a gene signature associated with DUSP11 deficiency in T cells composed of the top 100 most significantly upregulated genes (Table S3).

### DUSP11 downregulation and subsequent transcriptional response is observed in HIV-1 infected patients

To better characterize the relevance of DUSP11 in HIV-1 infection, we infected primary cells from 3 healthy donors with HIV-GFP, FACS-sorted the productively infected fraction (GFP+) and performed total RNAseq on GFP+ and GFP-fractions. We computed the DUSP11^-/-^ signature established in Jurkat (Table S3) on HIV-1 infected cells (GFP+) and compared it to non-(productively) infected cells (GFP-). Importantly, the DUSP11^-/-^ signature was sufficient to cluster both cell populations, independently of donor origin (Fig. 4A). We next interrogated the presence of a transcriptional signature similar to that caused by DUSP11 deficiency in HIV-1 patients. In a cohort of HIV-1 positive (HIV+) patients, for which PBMCs were collected prior to and 6 months after antiretroviral treatment (ART) (Table S4)(*25*), we performed total RNAseq on CD4+ T cells from HIV+ patients prior to ART and also found that the DUSP11^-/-^ gene signature was sufficient to cluster HIV+ patients from non-infected controls (Fig. 4B). Moreover, DUSP11 protein levels were significantly increased in 5/6 patients after antiretroviral treatment (Fig. 4C), indicating a HIV-1-mediated downregulation of DUSP11 before ART. Altogether, these results suggest that HIV-1 VPR actively alters the innate immune transcriptional response of HIV-1 patients through the direct targeting of DUSP11, which functions as a mediator of immunostimulatory properties of endogenous RNAs.

**Figure 4.**
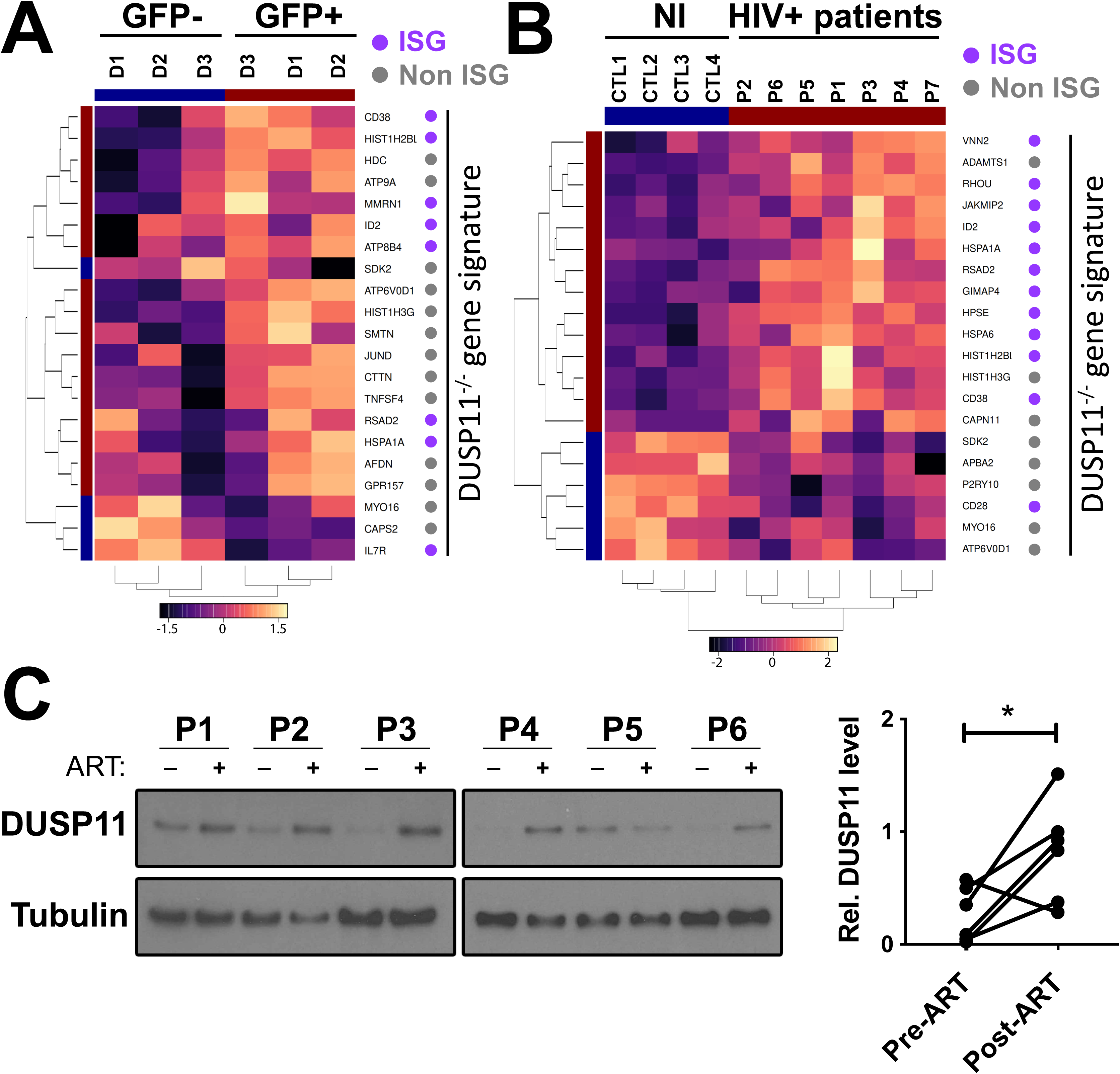
DUSP11 downregulation and subsequent transcriptional response is observed in HIV-1 infected patients. **A.** Hierarchical clustering and heatmap based on genes from DUSP11^-/-^ gene signature, differentially expressed between primary cells from 3 different donors, productively (GFP+), or non-productively (GFP-) infected with HIV-GFP. **B.** Hierarchical clustering and heatmap based on genes from DUSP11^-/-^ gene signature, differentially expressed between CD4 T cells from non-infected patients and acutely infected non-treated HIV+ patients. **C.** Western blot (left) and relative quantification (right) showing DUSP11 level in CD4 T cells from HIV+ patients prior and after anti-retroviral treatment. Paired t-test *p<0.05.

## Discussion

Here we describe the contribution of endogenous RNA sensing by RIG-I to innate immune responses elicited during RNA virus infection. In principle, any RNA transcribed by RNA Pol3 may have the ability to trigger RIG-I dependent immune responses, at least transiently, because they initially contain 5’-PPP terminal regions upon initiation of transcription. Indeed, a few of the Pol3-dependent RNAs, including RN7SL, RNA5S and vault-RNAs, have been shown thus far to trigger immune responses in different settings (*1, 3, 6, 10, 11, 26, 27*). In this work, we specifically identified the repeat family of Y-RNAs, and in particular RNY4, as a model of endogenous RNAs whose unique structure confers a previously unknown function as RIG-I agonists. We observed a contribution of Y-RNAs in RLR signaling during infections by MV and DV-4, both RNA viruses replicating in the cytoplasm and producing RLR-specific viral RNA ligands (*15, 16*). MV and DV-4 are acute viral infections where the speed of host response is critical to halt and ultimately clear viral replication. We speculate that the viral-mimicking structure of these endogenous RLRs ligands licenses them to act as innate immune guardians that prime immune responses at the onset of cell infection.

We observed the same contribution of Y-RNAs during infection by HIV-1 where we failed to detect any ligand of viral origin to either RIG-I, or MDA5, or LGP2. Further, we identified the cellular triphosphatase DUSP11 as a key immune modulator that prevents unwarranted sensing of cellular RNAs in heathy cells. Importantly, we show that HIV-1 has evolved mechanisms to manipulate and subvert this process through a targeted, VPR-dependent, degradation of DUSP11, leading to the subsequent activation of RIG-I by cellular RNAs (Fig. S4). While we cannot exclude that other functions associated with DUSP11, such as the maturation of miRNA (*20, 21*), might explain the selection forces leading to its targeting by HIV-1, we can also speculate that RLR activation by Y-RNAs may act as a rapid response mechanism for hosts to detect viruses that degrade DUSP11. Importantly, during the chronic phase of HIV-1 infection, higher levels of IFN-I signaling correlate with sustained levels of inflammation, immune exhaustion, CD4 T cell depletion and disease progression (*28*). In fact, chronic IFN-I signaling is considered by many as central to HIV-1 pathogenesis, to the point where the use of IFN-I blockade treatment is discussed as a supplement during ART (*29*). In this context, it will be important to determine whether, and to which extent, the immune activation subsequent to DUSP11 downregulation participates to the disruption of immune system homeostasis observed in HIV-1-infected patients. Finally, our results emphasize the contribution of PRRs in sensing not only microbial ligands but also self-derived ligands. In the context of host-pathogen interactions, these endogenous ligands, possibly owing to their molecular mimicry of pathogen-associated features, constitute a new class of immunostimulatory molecules that provide the host the unique advantage to control both their potency and accessibility to innate sensors.

## MATERIAL AND METHODS

### Cells lines

HEK293 (293, ATCC CRL-1573) and HEK293T (293T, ATCC CRL-3216) cells were maintained in DMEM-Glutamax supplemented with 10% heat-inactivated fetal calf serum (FCS, ThermoFisher Scientific) and Penicillin-Streptomycin (PS, Life Technologies). Jurkat T cells (Gift from Brown laboratory, Mount Sinai), MT4C5 cells (a derivate MT4 cells expressing CCR5) were used for co-culture with 293-4×4 and were cultured as described in (*1*). Primary T cells were maintained in RPMI supplemented with 5% pooled human serum (Gimini Bio-products) and HEPES buffer, non-essential amino acids, PS and L-glutamine (all Life Technologies). One-STrEP-tagged RLRs (ST-RLR: ST-RIG-I, ST-MDA5, ST-LGP2), CHERRY (ST-CH) cells (described in (*2*)) and STING-37 cell line corresponded to HEK293 cells stably transfected with an ISRE-luciferase reporter gene (described in (*3*)) were maintained in DMEM-Glutamax supplemented with 10% heat-inactivated FCS and 100 U/ml/100 mg/ml of PS and G418 (SIGMA) at 400 μg/ml. HAP1 RIG-I ko., LGP2 ko., MDA5 ko., MAVS ko. And control cell lines were purchased from Horizon Discovery (cat# HZGHC001441c001, HZGHC002927c011, HZGHC001448c012, HZGHC001456c011 and C631, respectively) and maintained in Iscove’s Modified Dulbecco’s Medium (ThermoFisher Scientific) with 10% FCS and PS. In order to generate ST-RLR cells susceptible to HIV-1 infection, they were transduced with lentiviral vectors encoding HIV-1 receptor (CD4) and co-receptor (CXCR4). After transduction, cells were sorted for the high level of expression of both CD4 and CXCR4 receptors. These cell lines were assigned ST-RLR-4X4. Stable cell line (assigned ST-CH-4X4) expressing the Cherry protein instead of tagged RLRs was generated and used as a negative control to allow subtraction of non-specific RNA binding.

### Generation of CRISPR-edited cell lines

293T knock out cell lines were generated by cotransfection (lipofectamine 2000, Invitrogen) of CRISPR–Cas9-expressing knockout plasmids (MAVS: sc-400769-ko-2, RIG-I: sc-400812, both Santa Cruz) and Homology Directed Repair plasmids containing puromycin resistance gene (MAVS: sc-400769-HDR-2, RIG-I: sc-400812-HDR, both Santa Cruz). The knockout plasmids are a mixture of three plasmids, each carrying a different guide RNA specific for the target gene, as well as the Cas-and GFP-coding regions. 72h after transfection cells were treated with puromycin (Invivogen, 1μg/ml) for 1 week. 293 and 2934×4 knock-out clones were generated by transfection (lipofectamine 2000, Invitrogen) of CRISPR–Cas9-expressing knockout plasmids (MAVS, sc-400769-ko-2 and control, sc-418922, both Santa Cruz). Jurkat knock-out clones were generated by electroporation (*Neon* Transfection System, Thermo Fisher Scientific) of CRISPR-Cas9-expressing knock-out plasmid (DUSP11, sc-408162; control, sc-418922, both Santa Cruz). 48h after transfection (293 & 2934×4) or electroporation (Jurkat), GFP^+^ cells were selected by cell sorting, and single clones were isolated in 96-well plates then cultured for 2 weeks. Depletion of target proteins was verified by western blotting.

### Affinity chromatography of RLR-RNP complexes and RNA extraction

ST-RLR cells were infected with MV (MOI=1) or DV-4 (MOI=0.5) for 24 hr or left uninfected (NI). In the case of HIV-1, ST-RLR4X4 (tagged) cells were co-cultivated with HIV-1-infected cells MT4C5 cells as described in (*1*). Briefly, 5.10^7^ MT4C5 (donor cells) were exposed 150ng (equivalent p24) of HIV-1 NL4.3 for 2hr at 37°C. After washing the virus, the cells were grown for 48h. The infection was monitored by flow cytometry analysis by intracellular Gag staining. Infection was then performed via coculture of ST-RLR4X4 cells and MT4C5 cells at a donor:target cell ratio of 1:1. 24 h post infection, cells were lysed and affinity purification of ST-tagged proteins and RNA extraction was performed as described in (*2, 4*).

### Isolation of primary cells

#### Isolation of T cells from healthy donors

PBMCs were prepared by Ficoll (GE Healthcare) gradient centrifugation from buffy coats received from New York blood center (Long island city, NY, USA). Buffy coats were diluted in 1:2 ratios (v/v) with PBS and 30 ml of the diluted buffy coats were loaded on 15 ml Ficoll in 50 ml falcon tubes. The tubes were centrifuged for 25 minutes at 2000 rpm at low acceleration and break. Mononuclear cells were collected and pooled from the tubes and washed twice by centrifuging. CD4+ T cell isolation was performed through beads-mediated negative selection (EasySep Human CD4+ T Cell Isolation Kit, Stemcell Technologies) and CD4+ cell purity was assessed by flow cytometry.

#### Isolation of T cells from HIV-1 cohort patients

Frozen PBMC from a cohort of intravenous drug using HIV-1+ patients (described in(*5*)) were thawed at 37°C and dead cells were removed through Annexin V beads-mediated selection (EasySep Dead Cell Removal Annexin V Kit, Stemcell technology). CD4+ T cells were further isolated through beads-mediated negative selection (EasySep Human CD4+ T cell Isolation Kit, Stemcell technology) and resuspended in Trizol (Ambion) (a small fraction was resuspended in PBS to check viability and purity by flow-cytometry). After addition of chloroform and phase separation, the top aqueous phase was used to subsequently isolate cellular RNA and the bottom organic phase was used to purify cellular proteins. Metadata of the cohort patients is listed in Table S5.

### *In vitro* transcription

*In vitro* transcribed (IVT) RNAs were generated using T7 RiboMAX express large-scale RNA production system (Promega), using oligoDNA containing the sequence of interest behind a T7 promoter. For full-length Y-RNAs, dsDNAs covering the entire sequence were used as templates. For RNY4 substructures fragments, a single oligo corresponding to the specific cDNA sequence of interest was annealed to another sense oligo containing the T7 promoter sequence, generating a DNA molecule as template where only the T7 promoter sequence was dsDNA. Sequences of DNA strands are listed in table S6 below, with T7 sequence in bold. T7 reaction mixed where then treated with DNAse to remove DNA template and purified using RNeasy kit (Qiagen). When specified, IVT RNAs were additionally treated with Antarctic phosphatase (NEB) to remove their 5’-PPP moieties then repurified. When indicated, RNAs were also generated from *in vitro* transcription of a modified p2RZ plasmid (Addgene #27664) where RNY4 or RNY4ds3 sequences were cloned between T7 promoter and HDV Ribozyme sequences.

### Luciferase-based Reporter Assay

*ISRE & IFN-*β *promoter reporter assays.* 293T cells, 293, 293-4×4 or HAP1 cells were seeded in 24-well plates. 24h later, reporter plasmids p-ISRE-Fluc (containing 5 ISRE promoter sequences upstream of the Firefly luciferase gene) or pIFNβ-Fluc (containing the IFN-β promoter upstream of the Firefly luciferase gene), and pTK-Rluc (containing a thymidine kinase promoter upstream of the Renilla luciferase gene) were transfected at a concentration of 100 ng/ml and 10 ng/ml, respectively. For experiments measuring responses to *in vitro* transcribed RNAs, plasmids were transfected together with 30 ng/ml of RNA of interest using Lipofectamine 2000 (Invitrogen). For experiments measuring responses to virus infection, cells were infected 24h later with MV (MOI 1) or DV-4 (MOI 0.5), or co-cultured with MT4C5 infected with HIV-1 at a donor:target cell ratio of 1:1. 24h later cells were lysed (Passive Lysis buffer, Promega) and Firefly luciferase and Renilla Luciferase activities were measured using Dual Luciferase Reporter Assay system (Promega). Renilla values were used as transfection normalization control. Low molecular weight and high molecular weight poly(I:C) (Invivogen) were used as positive control of activation at a concentration of 5 and 30 ng/ml, respectively.

*STING-37 assay.* STING-37 cells, corresponding to HEK293 cells stably transfected with the ISRE-luciferase reporter gene (described in (*3*)) were plated in 24-well plates. 24h later, cells were transfected with 5-20 ng/well of *in vitro* transcribed RNA using lipofectamine 2000 (Invitrogen). 24h after transfection, cells were lysed (Passive Lysis buffer, Promega) and Firefly luciferase activity was measured using the Bright-Glo Luciferase Assay System (Promega).

### Infection with virus / transduction with vector

HIV_NL4.3_-GFP, HIV_NL4.3_WT and HIV_NL4.3_ΔVPR (described in(*6, 7*)) were freshly produced through transfection (Polyjet, Signagen) of 293T cells with plasmids coding for the full-length viral genomes. 3^rd^ generation lentiviral vectors coding for ovalbumin (control) or VPR *WT* or a Q65R VPR mutant were produced through co-transfection of 293T cells with 4 plasmids coding for Gag/Pol (Addgene #12251), Rev (Addgene #12253), VSVg (Addgene #12259), or one of the transgenes mentioned above (this article) at a molar ratio of 1/1/1/2.6. 48h after transfection, supernatants were collected, spun down and filtered to removed cellular debris and used to infect (HIV) or transduce (lentiviral vectors) T cell (Jurkat or Primary cells) by spinoculation in 96-wells plate (1,200 x g, 90 min at 16°C) with polybrene (4 μg/ml, EMD Millipore). 6h after infection / transduction, supernatants were replaced with fresh medium. In the case of primary cells, CD4 T cells were activated using PHA-L (2 μg/ml) for 72h prior to infection.

When specified, HIV-GFP productively-infected cells were sorted based on the GFP expression on a FACS aria (BD biosciences) with biosafety cabinet facility. The cells were stained for CD3 and CD4 expression to identify CD4 T cell populations and CD8, CD14, CD19 and CD56 to identify other contaminating immune cell populations. Viable cells were discriminated with a viability dye (blue fluorescent dye, Thermofischer). CD3^+^CD4^+/dim^ GFP^+^ cells were sorted as infected and the CD3^+^CD4^+^GFP^-^ cells were sorted as non-infected to the collection tube and used for RNA and protein isolation. The MV-Schwarz vaccine strain (GenBank accession no. AF266291.1) has been previously described^(*8*)^. DV-4 strain Dominica (AF326825)^(*9*)^ was obtained from the Centro de Ingeniería Genética y Biotecnología (CIGB), Cuba.

### Differential 5’-PPP RNA digestion

1ug of total cellular RNA was treated with RNA 5’ polyphosphatase (enzyme that converts 5’-triphosphorylated RNA into 5’-monophosphorylated RNA, Lucigen) for 30 min at 37°C or mock-treated. RNAs were then purified and treated with Terminator™ 5’-Phosphate-Dependent Exonuclease (processive 5’ to 3’ riboexonuclease that specifically digests RNA with 5’-monophosphate ends, Lucigen) for 90 min at 30°C or mock-treated. Resulting RNAs were then purified and processed for qPCR analysis.

### RNA purification

Total cellular RNA was extracted using Trizol-Chloroform phase separation (Ambion) followed by RNA purification from the aqueous phase using a modified version of the Zymo RNA Clean and Concentrator (Zymo Research), through the addition of 2x volumes of ethanol to increase the retention of small RNA species. RNA was then subjected to DNAse digestion (TURBO DNA-free Kit, Invitrogen) then purified again using Zymo RNA Clean and Concentrator and finally resuspended in DPEC-treated water.

### Quantitative PCR

1ug of total RNA was converted to cDNA using RNA to cDNA EcoDry Premix Double Primed (Clontech) and resulting cDNA was diluted 10X in water. For differential enzymatic digestion analysis, qPCR reactions were carried out in 10 μl reaction volumes with 5 μl of TaqMan Fast Advanced Master Mix (Thermo Fisher Scientific), 2 μl of Primers/Probe mix and 3 μl of each cDNA sample. For all other analysis, qPCR reactions were carried out in 10μl reaction volume with 5 μl iTaq Universal SYBR Green Supermix (Bio-rad), 2 μl Primer mix and 3 μl of each cDNA sample. The qPCR reactions were run using a CFX384 Touch Real-Time PCR Detection System (Bio-Rad) in clear wells plates. Targets amplification were quantified using the ΔΔ*C*t method relative to β-actin. The list of the primers used in this study is provided in Table S7.

### Western Blotting

Whole-cell lysates were resuspended in Laemmi sample buffer (Bio-rad) completed with 10% *β*-*mercaptoethanol* and heated for 5 minutes at 95°C. 10-15ul of lysates were loaded onto 10% or 4-12% mini-protean TGX gels (Bio-rad) and the gel was run in Tris/Glycine/SDS buffer (Bio-rad). Proteins were transferred to 0.45 um PVDF membranes (Immobilon) in Tris/Glycine buffer (Bio-rad) supplemented with 20% methanol. Membranes were blocked in Tris-buffered saline (Bio-rad) plus 0.1% Tween 20 (Fisher) (TBS-T) containing 5% non-fat dry milk for 30 min at room temperature followed by overnight incubation with primary antibody at 4 °C. Membranes were then washed with TBS-T and incubated with HRP-conjugated secondary antibodies for 3 h at room temperature. Membranes were then washed and HRP was activated with ECL Plus Western Blotting Substrate (Pierce) for 5 minutes before being exposed to *CL-Xposure Film* (Thermo scientific). Relative HRP signals were quantified using image Lab software (Bio-rad), relative to GAPDH or Tubulin controls.

### Antibodies

#### Western Blot and Protein purification

The following antibodies were used: HIV-1 p24 (mouse monoclonal, Abcam, clone 39/5.4A, #ab9071); MAVS (rabbit polyclonal, Cell Signaling Technology - CST, #3993); DUSP11 (rabbit polyclonal, Proteintech, #10204-2-AP); α-tubulin (mouse monoclonal, Proteintech, clone 1E4C11, #66031-1-Ig); HIV-1 gp120 (sheep polyclonal, gift from Benjamin Chen laboratory); GFP (rabbit monoclonal, CST, clone D5.1, #2956); GAPDH (rabbit monoclonal, CST, clone 14C10, #2118); VPR (rabbit polyclonal, Proteintech, # 51143-1-AP), StrEP-Tag (mouse monoclonal, Qiagen, #34850). Peroxidase-conjugated secondary antibodies against rabbit IgG (#7074) and mouse IgG (#7076) were purchased from CST.

#### Flow Cytometry

The following antibodies were used: anti-p24-FITC (mouse monoclonal, Beckman and Coulter, #KC-57); anti-CD3-Pacific Blue (mouse monoclonal, Biolegend #300330); anti-CD4-PE (mouse monoclonal, Biolegend, #300539); anti-CD19-APC (mouse monoclonal, Biolegend #302212); anti-CD14-APC (mouse monoclonal, Biolegend, #325608); anti-CD56-APC (mouse monoclonal, Biolegend, #318310); anti-CD8-PerCP/Cy5.5 (mouse monoclonal, Bdbiosciences, #341051).

### RNA-seq analysis of total and RLR-bound RNA

Protocols for NGS library preparation and NGS of total and RLR-bound RNA from MV, DV-4-infected cells have been described in (*2, 4*). Before RNA-seq analysis of total and RLR-bound RNA from HIV-1-and mock-infected cells, depletion of ribosomal RNA was done using the riboZero reagents included in the TruSeq stranded total RNA library prep kit (#20020596, Illumina). NGS libraries were generated following the manufacturer’s protocol. The indexed samples were multiplexed per 4 or 6 and sequenced on a HiSeq2500 sequencer (Illumina) to produce single-ends 65 bases reads, bearing strand specificity.

Bioinformatics analysis of NGS reads was performed using the RNA-seq pipeline from Sequana (*10*). Reads were cleaned of adapter sequences and low-quality sequences using cutadapt (*11*) version 1.11. Only sequences at least 25 nt in length were considered for further analysis. STAR version 2.5.0a (*12*) with default parameters, was used for alignment on the reference genome (Human genome hg19 from UCSC). Genes were counted using featureCounts version 1.4.6-p3(*13*) from Subreads package (parameters: -t gene -g ID -s 1). For statistical analysis of NGS data, count data were analyzed using R version 3.5.1 and the Bioconductor package DESeq2 version 1.20.0 (*14*). The normalization and dispersion estimation were performed with DESeq2 using the default parameters and statistical tests for differential expression were performed applying the independent filtering algorithm. For each virus, a generalized linear model including the replicate, beads and protein factors as well as the beads x protein interaction was set in order to test for the differential expression between the biological conditions. For each pairwise comparison, raw p-values were adjusted for multiple testing according to the Benjamini and Hochberg (BH) procedure (*15*) and genes with an adjusted p-value lower than 0.05 were considered differentially expressed. Bioinformatics analysis of NGS reads for viral reads was performed as described in (*4*). The MV-Schwarz vaccine strain (AF266291.1), DV-4 strain Dominica (AF326825) and HIV-1 NL4.3 (AF324493.2) were used as references.

### RNA seq analysis (others)

Protocols for NGS library preparation of Ribo-depleted total cellular RNA were performed by Genewiz. Raw Illumina reads were trimmed using trim_galore (Babraham bioinformatics) and cutadapt (*11*) version 1.18 with default settings. Reads were then mapped to the human genome (gencode annotation, build 38) and to repbase elements (release 20) using STAR aligner (*12*) version 2.6.1c. Aligned reads were assigned to genes using the featureCounts function of subread package using the annotation (*16*). This produced the raw read counts for each gene. Mapping and counting of the reads were done in two stages. First, reads were mapped to the human genome, and the counts were determined using the gencode annotation and the annotation derived from the repeatmasker output. Second, the reads which were not assigned to any feature in either gencode or repeatmasker annotation were re-aligned to the repeat consensus sequence (repbase). Counts obtained from repeatmasker and repbase corresponding to the same family were added together. Differential expression analysis was performed using DESeq2 package (*14*).

### Polymerase-III transcript annotation

Identification of regions of genome transcribed by polymerase-III (Pol3) remains a topic of active research (*17, 18*). In order to check if a given transcript is transcribed by Pol3 for our analysis, we created a curated list of tentative ncRNA transcripts that are likely transcribed by Pol3. In the list, we included all ncRNA families that are known to be transcribed by Pol3 (*18*) as well as ncRNA from hg19 genome assembly which overlap or are in proximity of an annotated Pol3 binding site or known Pol3 transcript, as based on curation of published datasets from Pol3 transcription studies from Refs (*19-23*).

### Modeling RNY4-like structure in transcripts

We follow the computational approach originally conceived in the work(*24*) to find Y-RNA homologs in genomes and used the RNAMotif tool to identify motifs that fold into a Y-RNA like structure. The RNAMotif software searches given RNA sequences for regions that are able to fold into a specified secondary structure. It identifies all regions in the RNA sequence capable of adopting the specified structure and calculates the free-energy contribution of the RNA region folded into the structure using the nearest-neighbor model for RNA(*25*). For our study, we use the following constraints for the “RNY4” structure search with RNAMotif. The motif is required to consist of (annotation illustrated as in Fig. S2B): stem (S1) of length ranging from 5 to 13 base pairs, a loop (L1) (with 0 to 2 unpaired nucleotides on 5’side and 1 to 3 nucleotides on 3’ end side), followed by a stem (S2) of length from 7 up to 11 base pairs, with a loop (L2) of size 6 to 17 unpaired nucleotides on the 5’ end side and 6 to 17 nucleotides on the 3’-end side, and a stem (S3) of length 7 to 17 base pairs, with a terminal hairpin loop of length ranging from 3 to 10 bases. In each stem, we allow up to 1 mismatch, and the stem base-pairs can contain both Watson-Crick and wobble base pairs. We only search for presence of the motif at the 5’ start of the transcripts, so for the evaluated sequences, we only consider the motif to be present if a possible RNY4 like structure (with assigned free energy by Turner model(*25*) smaller than 0) is detected by RNAMotif within 6 bases from the 5’-end of the transcript.

The sequence datasets evaluated for RNY4 motif presence were the following: human cDNA and non-coding RNA sequences (from hg38 reference genome assembly), complete genomes of positive-sense viruses with human host (Table S8, obtained from NCBI viral genome database(*26*)), and inserts in genome that were annotated as belonging to Y-RNA family in the repetitive DNA element database(*27*). For each sequence, we also constructed a scrambled sequence, which was obtained by randomly permuting all nucleotides in the respective sequence, so that the frequency distribution of respective nucleotides remains the same, but their order is random. The set of scrambled sequences was also used to search for RNY4-like motifs.

## Acknowledgements

NV would like to thank Alice Lepelley, Sarah Moyon and Miriam Merad for insightful discussions and suggestions. The authors would like to thank members of Tangy, Bhardwaj and Greenbaum labs for support and valuable discussions. We are grateful to Lisset Hermida and Gerardo Enrique Guillen Nieto (CIGB, Havana, Cuba) for providing DV-4 isolate and Namita Sajita and Benjamin Chen (Icahn school of Medicine at Mount Sinai, NYC, USA) for providing HIV-GFP clones and valuable technical help. VN and AK would like to thank Florence Guivel-benhassine (Virus and Immunity Laboratory of Institut Pasteur) and Commere Pierre-Henri (Plate-Forme de Cytométrie, IP) for technical support.

This work was supported by Agence Nationale de Recherches sur le SIDA et les hépatites virales 2012-1 and 2012-2 *HIV signature on RIG-I-like receptors* to AVK and FT; Agence Nationale pour la Recherche ANR-16-CE15-0025-01 *ViroStorm* to NJ and GB; Fondation pour la Recherche Medicale FDT20140931129 to RYSD, National Institutes of Health R01CA201189 and R01CA180913 to NB, R01AI081848 to NV, NB and BDG, 7R01AI081848-04 and 1P30CA196521-01 to B.D.G. B.D.G. was supported by a Stand Up To Cancer - National Science Foundation - Lustgarten Foundation Convergence Dream Team Grant sponsored by Stand Up to Cancer, the Lustgarten Foundation, the V Foundation and the National Science Foundation (NSF 1545935). B.D.G. is The Pershing Square Sohn Prize-Mark Foundation Fellow supported by funding from The Mark Foundation for Cancer Research.

---

**Table S5.** Metadata of HIV-1 cohort patient samples used in this study.

| Sample | Viral Load | CD4 Count |
| --- | --- | --- |
| **HIV negative** |  |  |
| CTL1 | NI |  |
| CTL2 | NI |  |
| CTL3 | NI |  |
| CTL4 | NI |  |
| **HIV positive** |  |  |
| P1 Pre | 1,060,000 | 460 |
| P1 Post | <50 | 866 |
| P2 Pre | 1,080,000 | 311 |
| P2 Post | <50 | 1213 |
| P3 Pre | 1,860,000 | 269 |
| P3 Post | <50 | 602 |
| P4 Pre | 120,000 | 355 |
| P4 Post | <50 | 758 |
| P5 Pre | 591,000 | 470 |
| P5 Post | <50 | 866 |
| P6 Pre | 525,000 | 503 |
| P6 Post | <50 | 491 |
| P7 Pre | 428,000 | 427 |
| P7 Post | <50 | 487 |

**Table S6.**
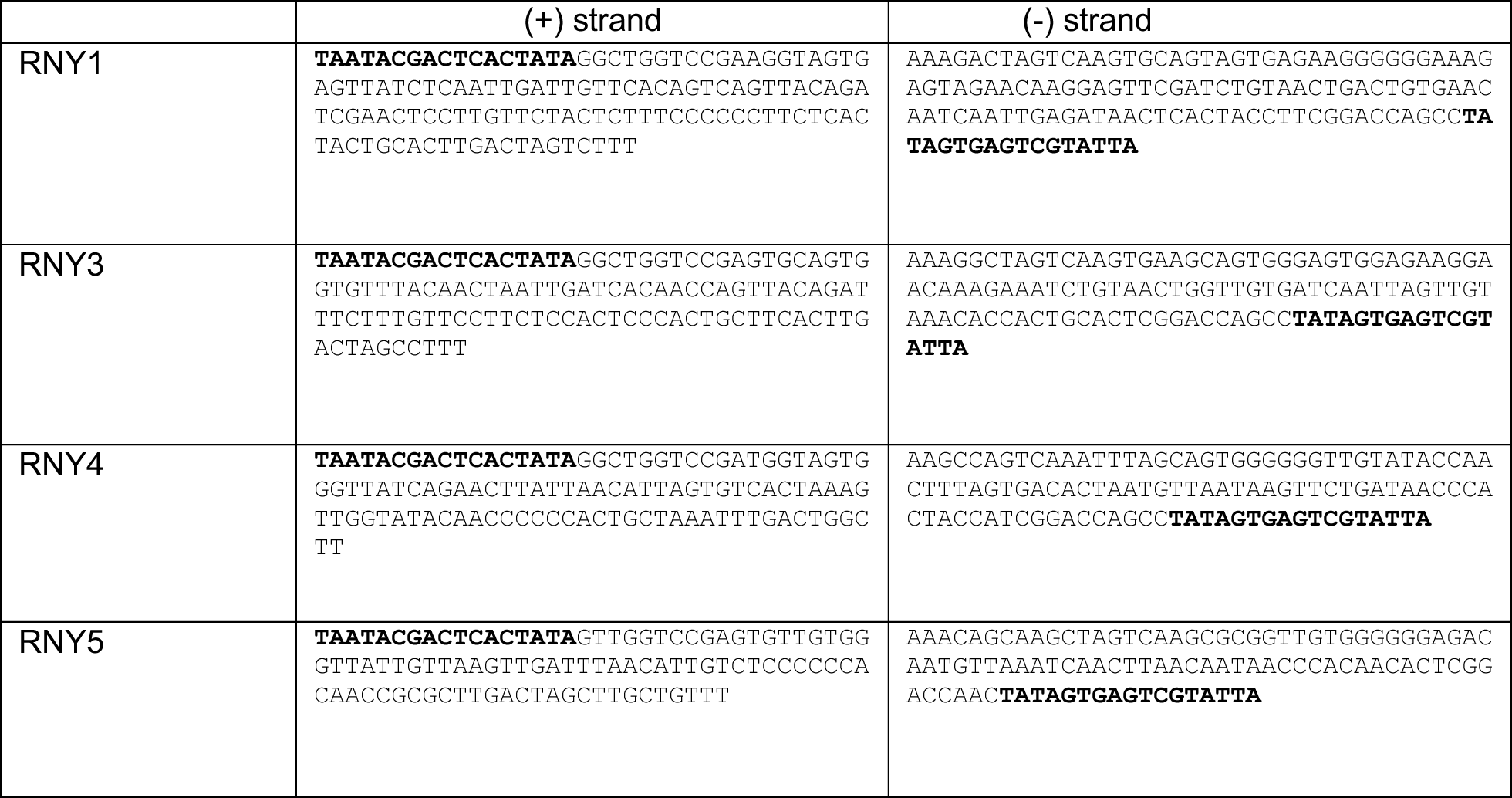
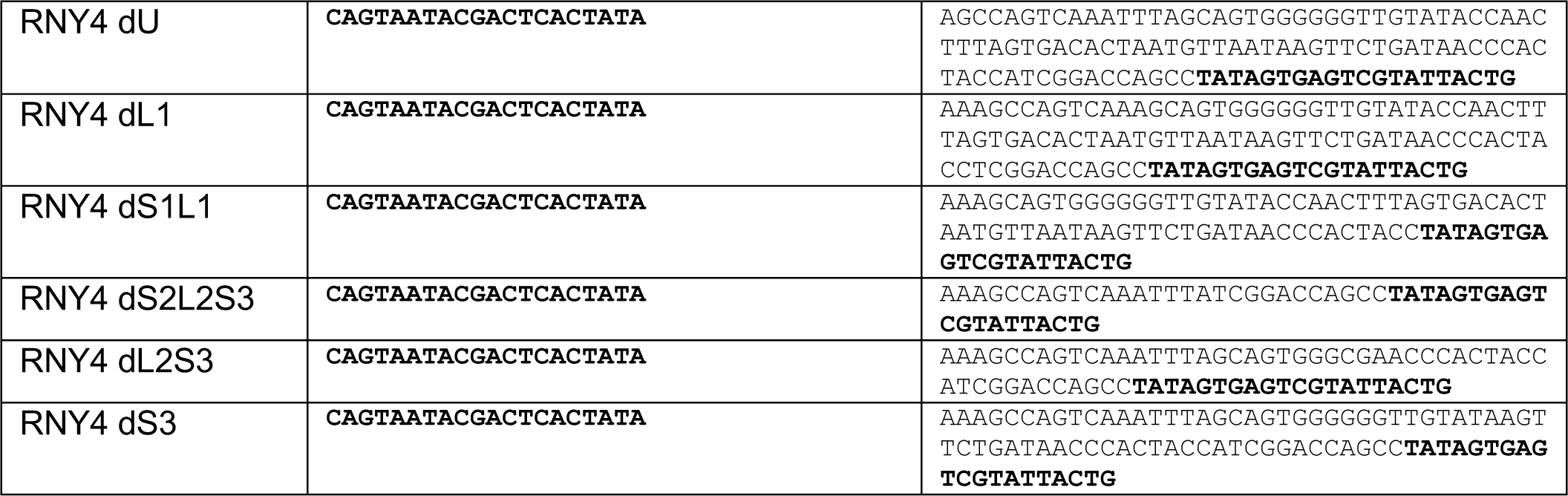
DNA template used for *in vitro* transcription

**TABLE S7.**
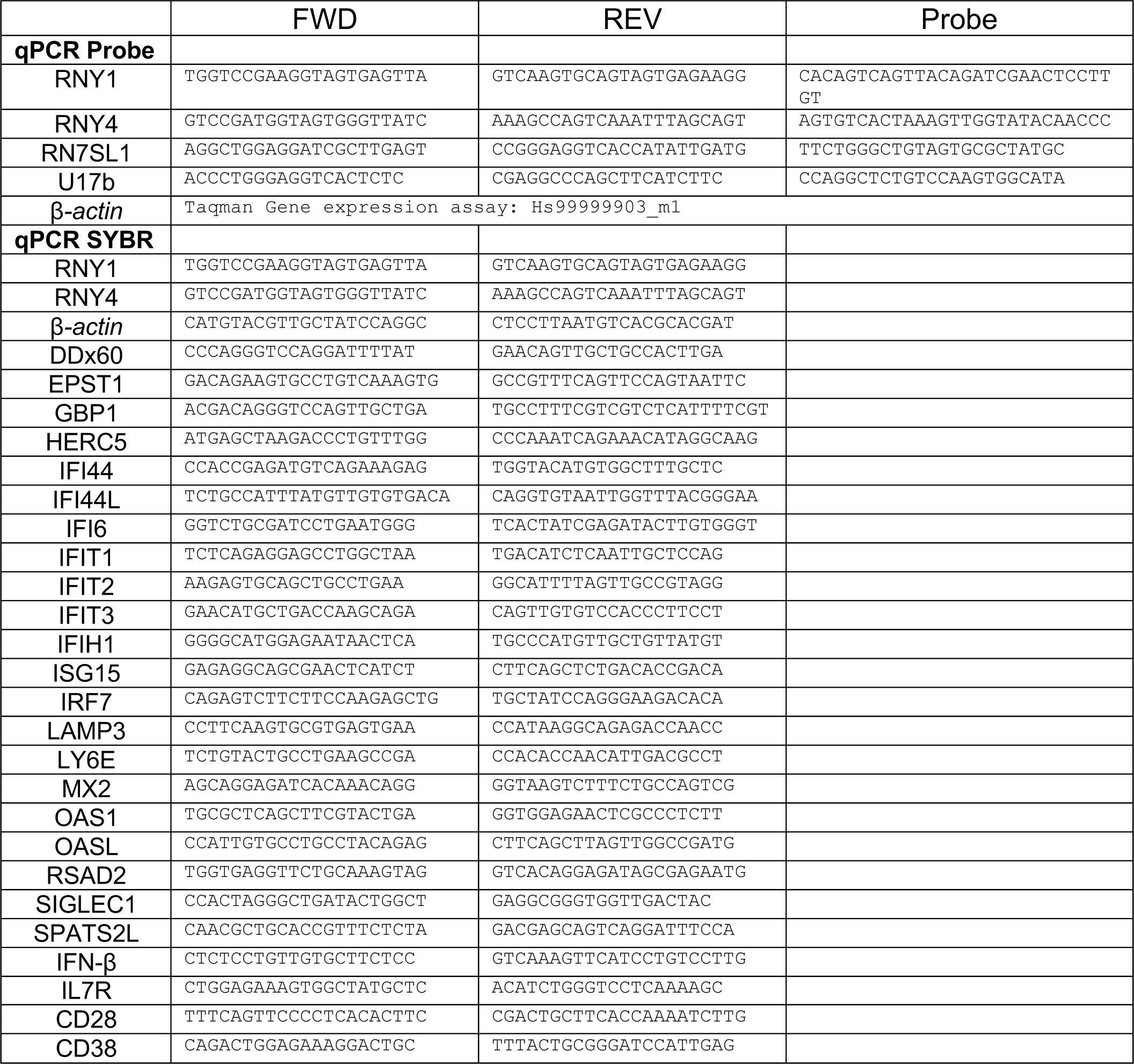
qPCR primers used in this study

**Supplementary Figure 1.**
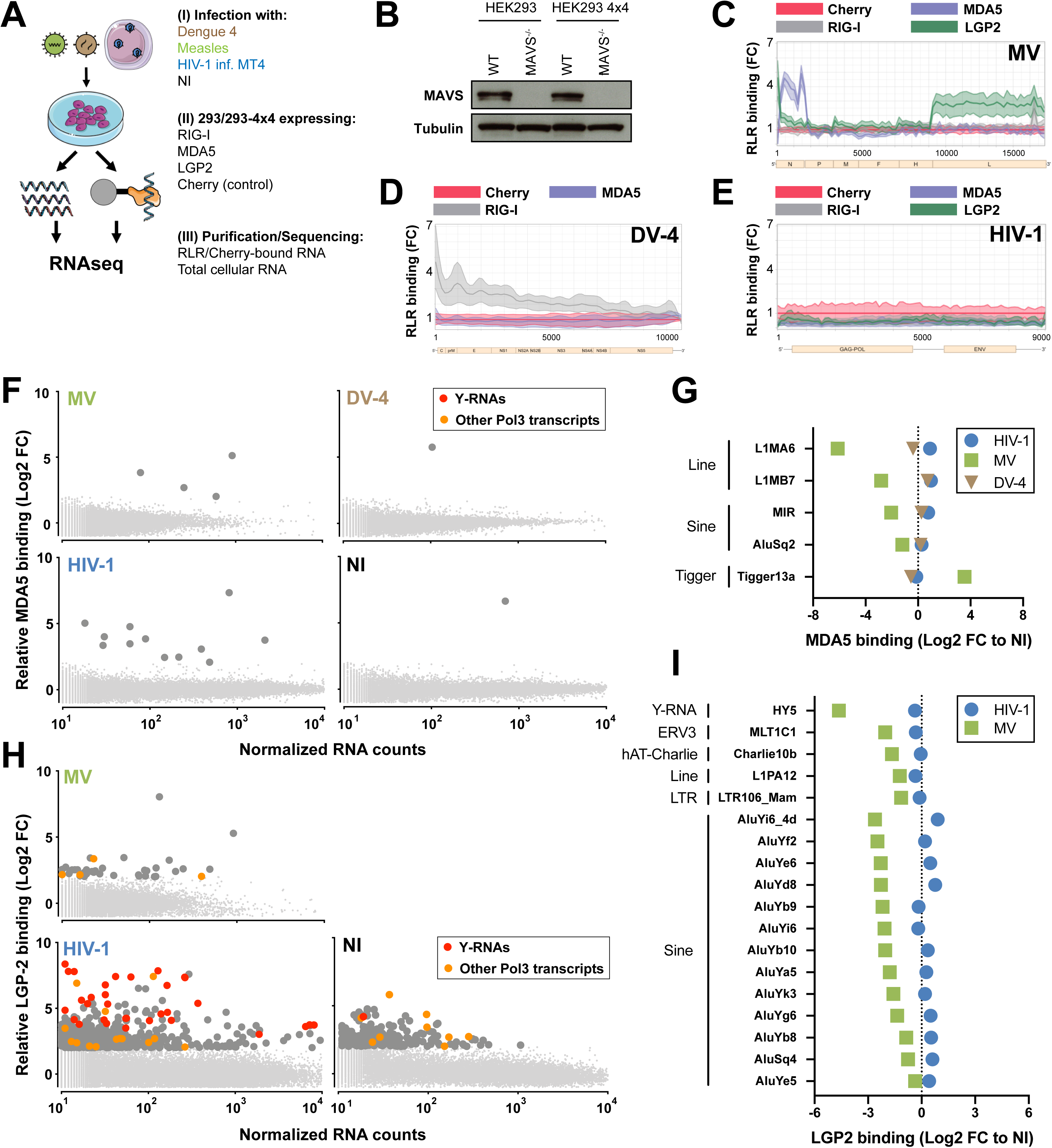
**A.** *Experimental approach* – 293 or 293-4×4 were engineered to stably expressed STrEP-tagged RLR receptors RIG-I, MDA5 or LGP2, or the non-RNA binding control Cherry. (I) 293 were infected with Measles Virus (MV) or Dengue Virus 4 (DV-4). 293-4×4 cells were cocultivated with HIV-1 infected MT4, non-infected cells were used as controls. (II) 24h after infection/coculture, cells were lysed, RLR receptors and Cherry protein were purified using STrEP-tag affinity and corresponding RNA fractions were isolated. (III) Total cellular RNA and RLR/Cherry-bound RNA fractions were subjected to total RNAseq and reads aligned on viral genomes (MV Schwarz strain; DV-4 Dominica; HIV-1 NL4.3) and human genome (HG38). **B.** Western Blot showing complete MAVS depletion in 293 and 293-4×4 MAVS^-/-^ cells. **C-E.** 24h post-infection with MV (**C**) or DV-4 (**D**), after coculture with HIV-1 infected MT4 (**E**), total cellular RNA and RLR/RNA purified complexes are subjected to strand-specific NGS analysis. Sequencing reads are mapped to MV (**C**) or DV-4 (**D**) or HIV-1 (**E**) genome after normalization based on total RNA samples. Differential enrichment analyses were performed between RLR/RNA and Cherry/RNA samples. The distribution of normalized read coverage matching each virus genome is represented along the (x-axis), and showing the fold enrichment on beads between RIG-I, MDA5 (**C**-**E**), LGP2 (**C**,**E**) compared to Cherry control. The curves were obtained from averaging read coverage of three independent experiments. The corresponding data were previously described in (15,16). **F.** 24h after infection with MV or DV-4, after coculture with HIV-1 infected MT4C5 or in NI control, sequencing reads were mapped to human genome. Differential enrichment analyses were performed between MDA5/RNA and Cherry/RNA samples. Genes are represented following their normalized count in cellular RNA (x-axis) and their fold enrichment (log2) to MDA5 compared to Cherry control (y-axis) from averaging three independent replicates. Genes that showed a log2(FC)>2 and adj-pval<0.05 are represented with larger dot size. Among these, RNA Pol3 transcripts are shown in orange and transcripts from Y-RNA families in red. **G.** Families of repeats RNA that show specific affinity to MDA5 compared to Cherry in at least one infected or NI condition, computed according to their relative enrichment compared to NI. **H.** 24h after infection with MV or after coculture with HIV-1 infected MT4 or in NI control, sequencing reads were mapped to human genome. Differential enrichment analyses were performed between LGP2/RNA and Cherry/RNA samples. Genes are represented following their normalized count in cellular RNA (x-axis) and their fold enrichment (log2) to LGP2 compared to Cherry control (y-axis) from averaging three independent replicates. Genes that showed a log2(FC)>2 and adj-pval<0.05 are represented with larger dot size. Among these, Pol3 transcripts are shown in orange and transcripts from Y-RNA families in red. **I.** Families of repeats RNA that show specific affinity to LGP2 compared to Cherry in at least one infected or NI condition, computed according to their relative enrichment compared to NI. **B.** Western Blot representative of n=3 independent experiments. **C-I.** Enrichment calculated from the mean of n=3 infection/RLR-purification/sequencing experiments.

**Supplementary Figure 2.**
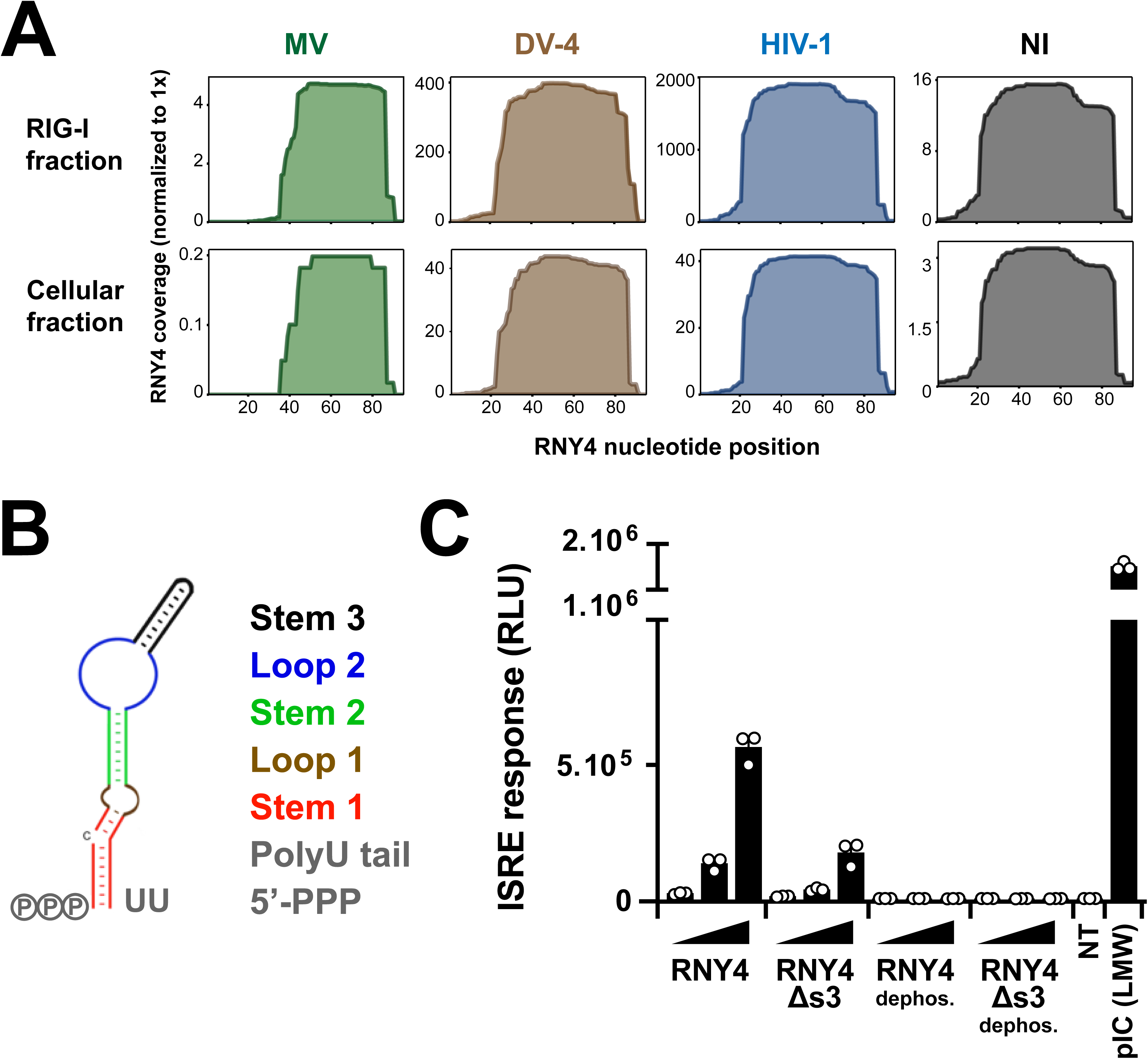
**A.** RNAseq normalized read coverage of RNY4 gene in RIG-I isolated fractions (top row) and cellular fractions (bottom row) in the different infection conditions. **B.** Schematic secondary structures (as described in 18) of RNY4 detailing the different molecular substructures analyzed in this work. **C.** Luciferase reporter activities showing ISRE-luciferase response from STING-37 cells transfected with 5-10-20ng/ml of different RNY4 subsets. RNY4 and RNY4dS3 RNAs were generated through *in vitro* transcription from a modified p2RZ plasmid where RNY4 or RNY4dS3 sequences are cloned downstream of a T7 promoter and upstream of a 3’ ribozyme sequence that generates discrete 3’ ends. Both RNAs were subsequently treated with Alkanine Phosphatase to remove their 5’-PPP moieties. **C.** Data representative of n=3 independent experiments. Bars show mean +/− s.e.m. of technical triplicates.

**Supplementary Figure 3.**
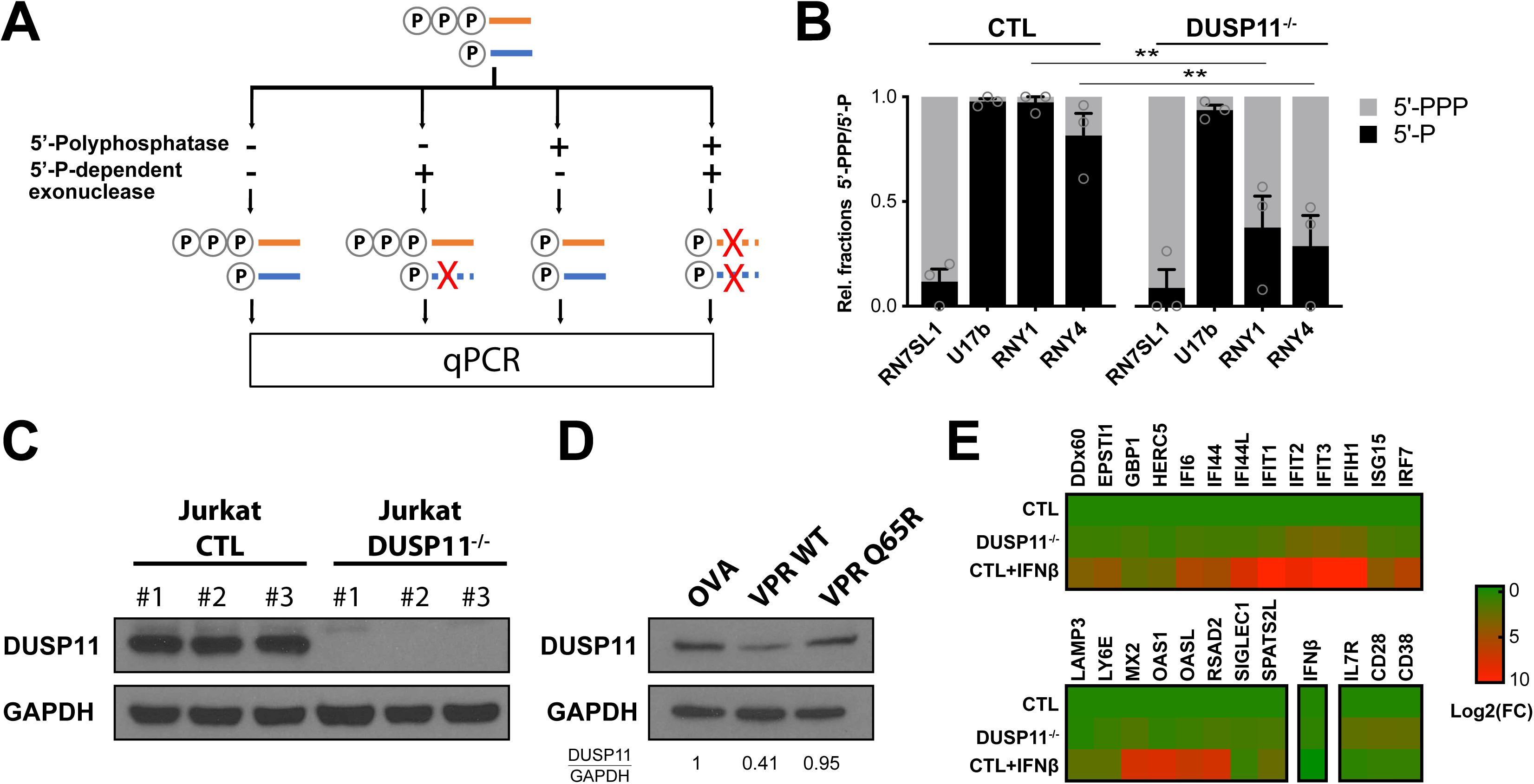
**A.** Summary schematic of differential enzymatic digestion. Total cellular RNAs were isolated and treated with either 5’-polyphosphatase, 5’-P-dependendent exonuclease or both consecutively. Resulting RNAs were purified, reverse-transcribed and their level measured by qPCR. RN7SL1 and U17b served as 5’-PPP and 5’-P controls, respectively. **B.** Ratio of 5’-PPP and 5’-P-bearing RNY1 and RNY4 in control or DUSP11^-/-^ Jurkat T cells. Relative 5’-PPP/5’-P RNA levels were determined through differential enzyme digestion followed by qPCR relative to β-actin mRNA. RN7SL1 and U17b served as 5’-PPP and 5’-P RNA controls, respectively. **C.** Western Blot showing complete DUSP11 depletion in Jurkat DUSP11^-/-^ clones. **D.** Western Blot showing DUSP11 depletion in Jurkat cells 72h after transduction with lentiviruses coding for ovalbumin control (OVA), HIV-1 VPR *WT* or a VPR(Q65R) mutant, defective for DCAF1 binding. **E.** Heatmap of qPCR values measuring expression level of a panel of IFN-I stimulated genes and markers of T cells activation (CD28, CD38, IL7R) in Jurkat control, DUSP11^-/-^, or control treated overnight with recombinant IFN-β. Expression levels are normalized to *β-actin* mRNA levels and to Jurkat control. **B.** Bars show mean +/− s.e.m. of 3 control and 3 DUSP11^-/-^ Jurkat clones. Student’s t-test **p<0.01 **C.** Western Blot representative of n=2 independent experiments. **D.** Western Blot representative of 3 independent experiments. Numbers at the bottom indicate semi-quantification of relative DUSP11/GAPDH levels normalized to control condition. **E**. Heat map show mean of 3 controla and 3 DUSP11^-/-^ Jurkat clones.

**Supplementary Figure 4.**
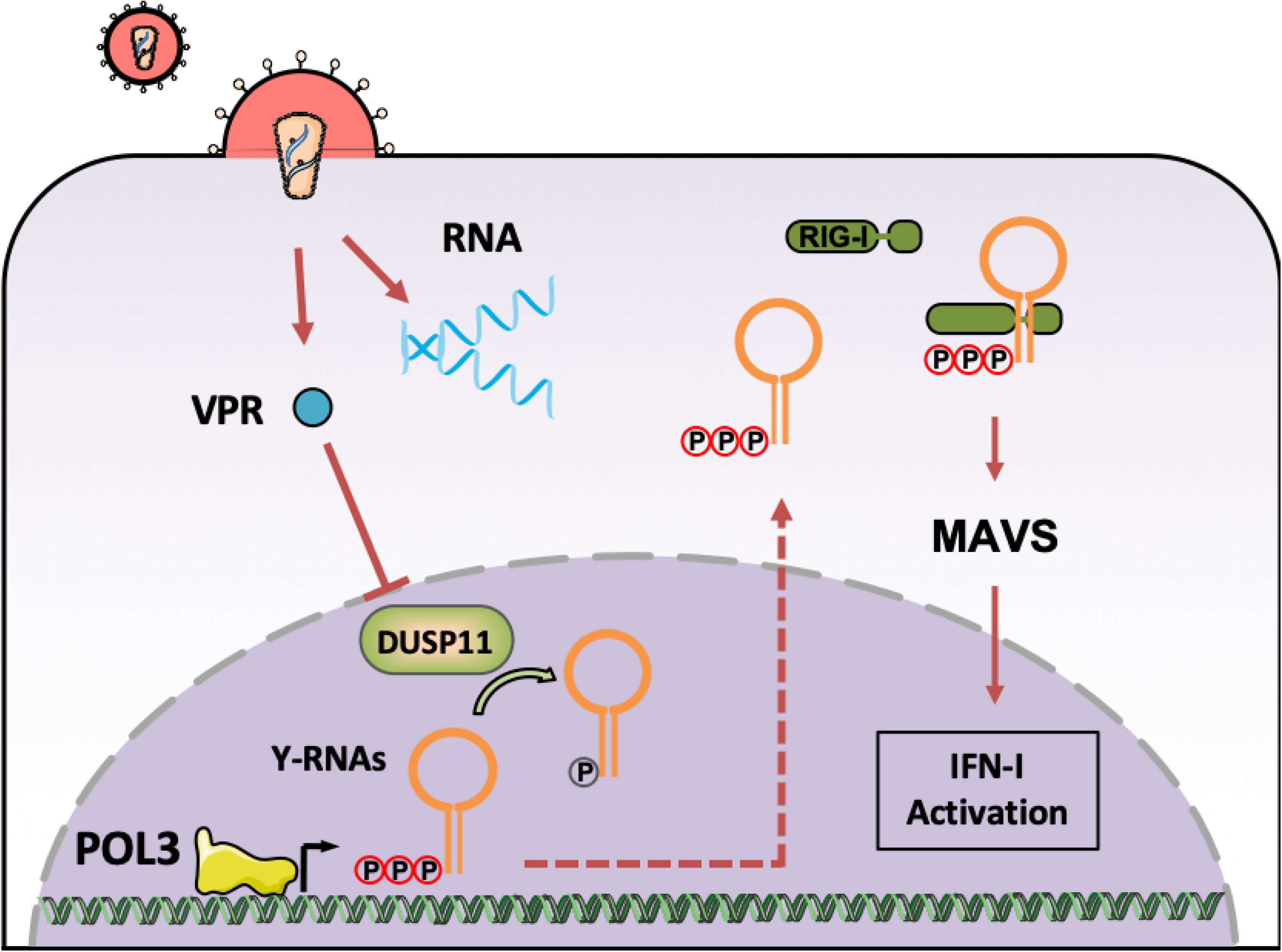
Schematic of the proposed mechanism for the contribution of Pol3-transcribed endogenous RNAs in the activation of RIG-I/MAVS during HIV-1 infection.

